# Age-induced methylmalonic acid accumulation promotes tumor progression and aggressiveness

**DOI:** 10.1101/2020.06.04.135087

**Authors:** Ana P. Gomes, Didem Ilter, Vivien Low, Jennifer E. Endress, Juan Fernández-García, Adam Rosenzweig, Tanya Schild, Dorien Broekaert, Adnan Ahmed, Melanie Planque, Ilaria Elia, Julie Han, Charles Kinzig, Edouard Mullarky, Anders P. Mutvei, John Asara, Rafael de Cabo, Lewis C. Cantley, Noah Dephoure, Sarah-Maria Fendt, John Blenis

**Affiliations:** Meyer Cancer Center, Weill Cornell Medicine, New York, NY, USA; Department of Pharmacology, Weill Cornell Medicine, New York, NY, USA; Laboratory of Cellular Metabolism and Metabolic Regulation, VIB-KU Leuven Center for Cancer Biology, VIB, Herestraat 49, 3000 Leuven, Belgium; Laboratory of Cellular Metabolism and Metabolic Regulation, Department of Oncology, KU Leuven and Leuven Cancer Institute (LKI), Herestraat 49, 3000 Leuven, Belgium; Weill Cornell Medicine/Rockefeller University/Sloan Kettering Tri-Institutional MD-PhD Program, New York, New York, USA; Department of Medicine, Weill Cornell Medicine, New York, NY, USA; Department of Medicine, Beth Israel Deaconess Medical Center and Harvard Medical School, Boston, MA, USA; Laboratory of Experimental Gerontology, National Institute on Aging, National Institutes of Health, Baltimore, MD, USA; Department of Biochemistry, Weill Cornell Medicine, New York, NY 10021, USA

## Abstract

From age 65 onwards, the risk of cancer incidence and associated mortality is substantially higher^1-3^. Nonetheless, our understanding of the complex relationship between age and cancer is still in its infancy^4^. For decades, the link has largely been attributed to increased exposure time to mutagens in older individuals. However, this view does not account for the well-established role of diet, exercise and small molecules that target the pace of metabolic aging^5-8^. Here, we show that metabolic alterations that occur with age can render a systemic environment favorable to progression and aggressiveness of tumors. Specifically, we show that methylmalonic acid (MMA), a by-product of propionate metabolism, is significantly up-regulated in the serum of older people, and functions as a mediator of tumor progression. We traced this to the induction of SOX4 and a consequent transcriptional reprogramming that can endow cancer cells with aggressive properties. Thus, accumulation of MMA represents a novel link between aging and cancer progression, implicating MMA as a novel therapeutic target for advanced carcinomas.

---

It has long been known that age is one of the main risk factors for cancer incidence ^9-11^. How it contributes to cancer progression, however, is not well understood^2,3,12^. Considering the growing body of evidence demonstrating that cancer cell-extrinsic factors are key in modulating tumor progression, we hypothesized that aging produces a systemic environment that supports tumor progression and development of aggressive properties. To test this hypothesis, we cultured A549 non-small cell lung cancer (NSCLC) and HCC1806 triple negative breast cancer (TNBC) cells in 10% human serum from 30 young (age ≤ 30) and 30 old (age ≥ 60) individuals with no diagnosed disease (Fig. 1a; Extended Data Table 1). While the majority of cells treated with young donor sera maintained their epithelial morphology (25 out of 30), cells treated with 25 out of the 30 old donor sera became mesenchymal, losing polarity and displaying a spindle-shaped morphology (Extended Data Fig. 1-3). These phenotypes were independent of donor ethnicity, and resembled epithelial-to-mesenchymal transition (EMT), a developmental process that during cancer pathogenesis can confer cells the cellular plasticity necessary to acquire pro-aggressive and pro-metastatic properties^13^. Cells cultured in the presence of sera from aged donors displayed a pronounced loss of the epithelial marker E-cadherin and gain of the mesenchymal markers fibronectin and vimentin^13^, in addition to increased expression of SERPINE1 and MMP2^14,15^ – proteins highly associated with aggressive phenotypes (Fig. 1b, Extended Data Fig. 4a, b). Moreover, the aged sera promoted resistance to two distinct and widely used chemotherapeutic drugs, carboplatin and paclitaxel (Fig. 1c, Extended Data Fig. 4c). Compelled by these observations, we questioned if the cells treated with the old donor sera would also display heightened metastatic potential. We treated MDA-MB-231 cells with sera from old or young donors before injecting them into the tail veins of athymic mice. The aged sera robustly potentiated the ability of the cells to colonize the lungs and form metastatic lesions, in contrast to the young sera (Fig. 1d, e). Together, our data show a role for systemic aging and age-induced circulatory factors in promoting the acquisition of aggressive properties of cancers.

**Fig. 1:**
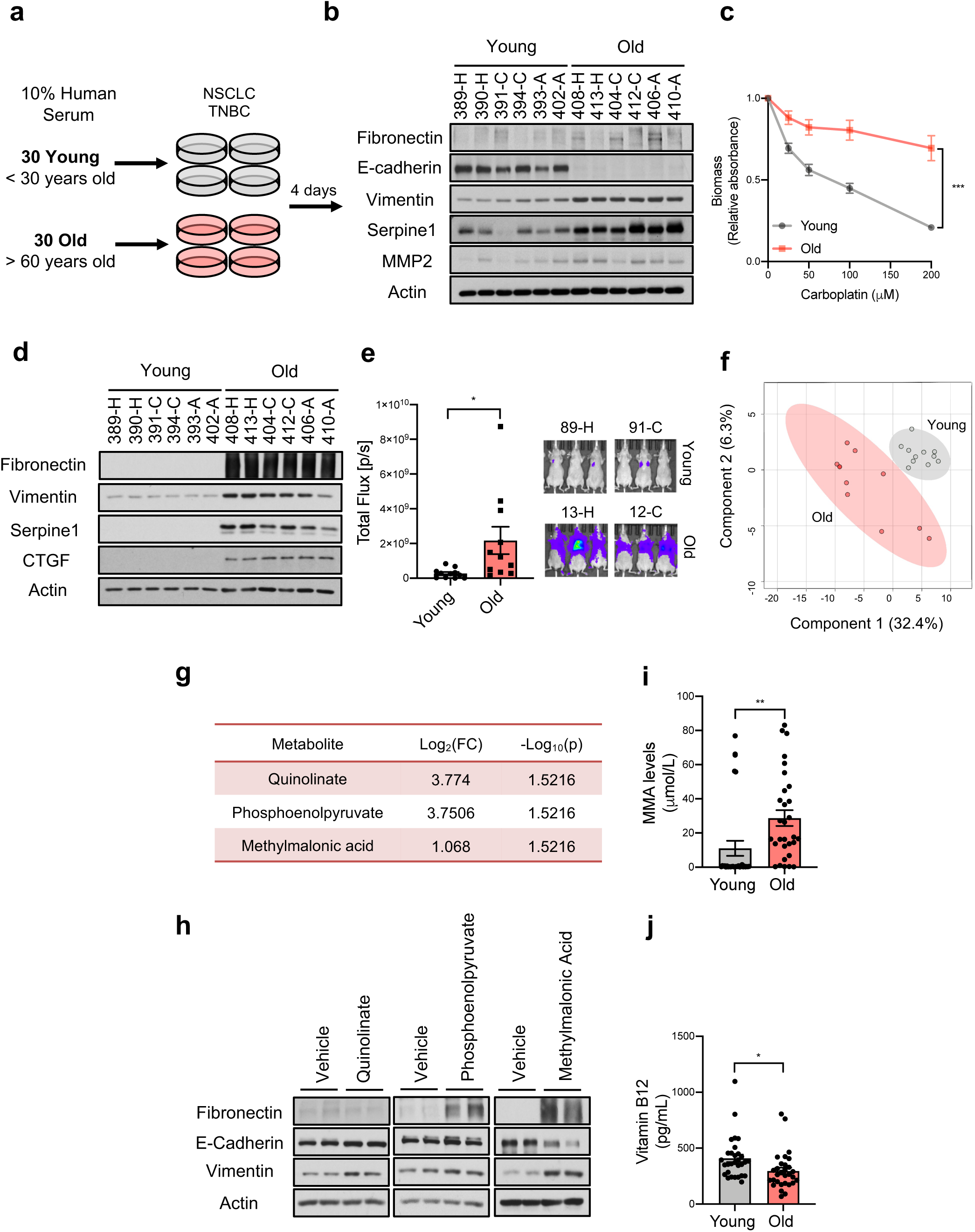
An age-induced circulatory factor promotes aggressive properties of cancer cells. **a**, Diagram showing experimental design (NSCLC: non-small cell lung cancer, TNBC: Triple negative breast cancer). **b**, Immunoblots for aggressiveness markers in A549 cells cultured for 4 days in 10% human serum from young and old donors; representative images, see Extended Data Fig. 4a (n=30). **c**, Resistance to carboplatin in A549 cells cultured for 4 days in 10% human serum (n=15). **d, e**, Metastatic properties of MDA-MB-231-luciferase cells cultured for 5 days in 10% human serum evaluated by immunoblots for aggressiveness markers (n=6) (**d**) and lung colonization assay (n=11) (**e**). **f**, PLSDA analysis of the targeted metabolomics of the sera (n=11). **g**, List of all metabolites that are increased at a statistically significant level in the sera of old donors (n=11). **h**, Immunoblots for aggressiveness markers in A549 cells treated with 5 mM methylmalonic acid (MMA), or 5 mM phosphoenolpyruvate or 5 mM quinolinate for 10 days; representative images (n=4). **i, j**, Concentrations of MMA (**i**) and vitamin B12 (**j**) in all human serum samples (n=30). For (**b, d**): A – African American; C – Caucasian; H – Hispanic. All values are expressed as mean ± SEM (*p<0.05, **p<0.01, ***p<0.001).

Pro-inflammatory factors play a key role in tumor progression and metastasis formation^16^, and are also known contributors of age-related diseases^17^. We first sought to test if a general pro-inflammatory signature was present in the old sera that could explain the induction of aggressive properties in cancer cells, but analysis of the circulatory proteome in these samples did not present such a signature (Extended Data Fig. 4d). As an alternative, we hypothesized that metabolic differences may be responsible for our observations, particularly considering the effectiveness of metabolic interventions such as diet, exercise and caloric restriction, in mitigating cancer susceptibility and outcome^1,18-20^. To identify potential candidates, we examined the metabolic compositions of the sera from the young and old donors (Fig. 1f). Out of the 179 circulatory metabolites detected by targeted metabolomics, only 10 were altered at a statistically significant level (Extended Data Table 1). A pronounced decline in circulating glutathione and spermidine levels was expected, considering their known roles in the aging process^21-24^. We also observed a substantial decline in serum levels of glutamine and α-ketoglutarate, supporting a previously suggested role for glutamine metabolism during aging^25^ (Extended Data Fig. 4e). Interestingly, only 3 metabolites were consistently increased in the sera from the aged donors: phosphoenolpyruvate, quinolinate and methylmalonic acid (MMA) (Fig.1g). To test if any of these 3 metabolites was responsible for inducing the pro-aggressive effects, we treated NSCLC cells with 5 mM of each metabolite. Only MMA induced a complete pro-aggressive EMT-like phenotype with a decline in E-cadherin and a concurrent increase in fibronectin and vimentin (Fig. 1h).

MMA, a dicarboxylic acid, is primarily a by-product of propionate metabolism. Propionyl-CoA, produced from catabolism of branched chain amino acids and odd chain fatty acids, yields succinyl-CoA in a vitamin B12-dependent manner to fuel the TCA cycle. Accumulation of MMA results from increased flux through and/or deregulation of the enzymes in this pathway and is a marker for a group of inborn metabolic diseases called methylmalonic acidemias, as well as of vitamin B12 deficiency^26^. Large-scale exploratory metabolomic experiments are notorious for their lack of sensitivity and quantitation^27^; therefore, to gain a deeper insight into the dynamics of MMA levels with age we determined the absolute concentration of this metabolite in the sera samples from all 60 donors. This deeper analysis revealed a more substantial increase in the levels of MMA in the sera of the old (15-80 μM) compared to the young donors (0.1-1.5 μM) (Fig. 1i). Moreover, in the case of the 10 outlier samples (5 old non-EMT-inducing and 5 young EMT-inducing donor sera) MMA levels consistently correlated with the phenotypes observed in cancer cells, supporting the idea that MMA is, at least in part, responsible for the observed age-related aggressive phenotypes (Extended Data Fig. 4f). Vitamin B12 levels are known to decline with age^28^, implicating vitamin B12 deficiency as a likely candidate for the age-induced accumulation of MMA. While measurements of vitamin B12 in the sera revealed a modest decline in old donors (Fig. 1j), this decline did not correlate with MMA levels in the outlier samples (Extended Data Fig. 4g). Although we cannot exclude vitamin B12 deficiency as a contributor to the accumulation of MMA with age, other factors such as deregulation of propionate metabolism in a major organ are also likely to be at play.

To better understand MMA’s pro-aggressive properties, we treated HCC1806, A549 and a breast epithelial (MCF-10A, a common model for EMT studies^29-31^) cell line with MMA. Concentrations of 1 mM and above were sufficient to induce an EMT-like phenotype and the expression of pro-aggressive proteins (MMP2, SERPINE1 and CTGF) (Fig. 2a, Extended Data Fig. 5a, b). Importantly, the pro-aggressive effects of MMA were specific, as different acids of similar structure and pKa did not induce the same phenotype (Extended Data Fig. 6a, b). MMA treatments also induced resistance to carboplatin and paclitaxel (Fig. 2b, Extended Data Fig. 5f-h), increased the cells’ migratory and invasive capacity (Fig. 2c, Extended Data Fig. 5c), and promoted stem-like properties, as shown by an upregulation of CD44 and a decline in CD24^32^ (Extended Data Fig. 5d, e). Treatment of MDA-MB-231 cells *in vitro* with MMA increased markers of aggressiveness (Extended Data Fig. 5i) and was sufficient to robustly increase their ability to colonize the lungs of athymic mice in a concentration-dependent manner (Fig. 2d). All together, these data support MMA as a promoter of pro-aggressive traits and as a contributor to the cellular plasticity required for tumor progression.

**Fig. 2:**
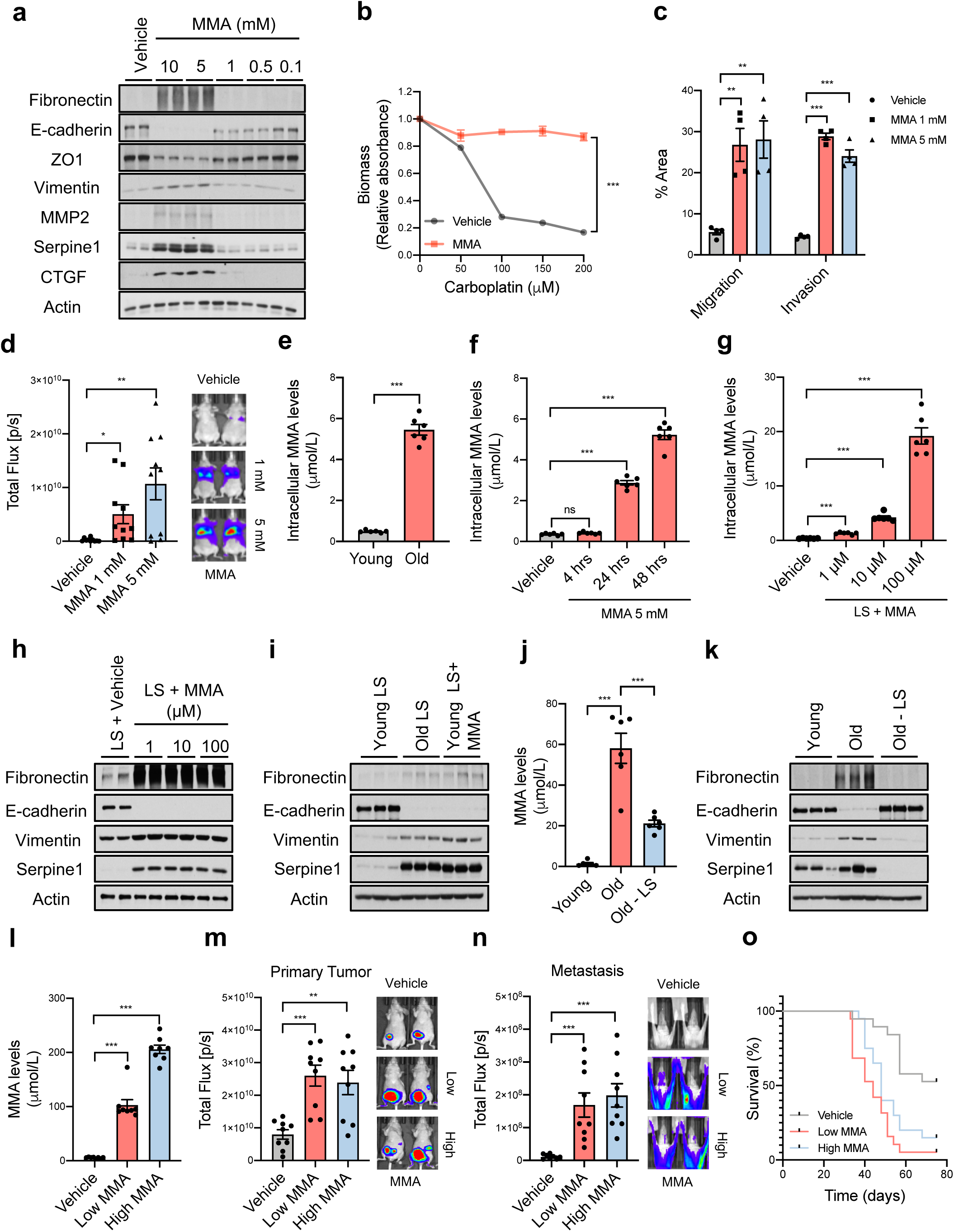
Methylmalonic acid induces aggressive traits. **a**, Immunoblots for aggressiveness markers in A549 cells treated with MMA for 10 days; representative images (n=4). **b**, Resistance to carboplatin in A549 cells treated with 5 mM of MMA for 10 days (n=4). **c**, Transwell migration/invasion assays of MCF-10A cells treated with MMA for 10 days (n=4). **d**, Lung colonization assay of MDA-MB-231-luciferase cells treated with MMA for 5 days (n=10). **e-g**, Intracellular MMA concentrations in A549 cells cultured with 10% human serum for 4 hr (**e**), with 5 mM MMA for indicated time periods (**f**), or with MMA-loaded FBS lipidic structures (LSs) for 4 hr (**g**) (n=6). **h, i**, Immunoblots for aggressiveness markers in A549 cells treated with MMA-loaded FBS LSs for 4 days (n=4) (**h**), or with LSs isolated from young/old serum, and MMA-loaded LSs isolated from young serum (10 μM MMA) (n=6) (**i**); representative images. **j**, MMA concentrations in serum from old donors after depletion of LSs compared to control human serum (n=6). **k**, Immunoblots for aggressiveness markers in A549 cells treated for 4 days with 10% human serum from old donors after depletion of LSs or from control donors (n=6); representative images. **l-n**, End-point serum MMA concentrations (n=8) (**l**), bioluminescence intensity of the primary tumors (n=9) (**m**), and metastases (n=9) (**n**) in mice that were xenografted with MDA-MB-231-luciferase cells and subcutaneously injected with MMA daily (n=8). **o**, Kaplan-Meier curve of mice xenografted with MDA-MB-231-luciferase cells and treated with MMA either subcutaneously or through drinking water. (n=19). All values are expressed as mean ± SEM (*p<0.05, **p<0.01, ***p<0.001).

Although MMA concentrations above 1 mM were required to potently induce aggressive traits in cancer cells *in vitro*, MMA concentrations measured in the sera from old donors were much lower (Fig. 1i, 2e). Further analysis demonstrated that the intracellular concentrations of MMA achieved within 4 hours of treatment with old sera or 5 mM MMA were substantially different (Fig. 2e, f). In fact, 5 mM MMA could only produce intracellular concentrations similar to the ones observed with 4-hour old sera treatment by 48 hours. In contrast, a more cell-permeable version of MMA (dimethyl MMA) could induce pro-metastatic effects at concentrations as low as 10-50 µM (Extended Data Fig. 6c, d), suggesting that the discrepancy in concentrations are due to a lower cell-permeability of added MMA compared to endogenous MMA in donor sera. To assess if another component of the sera could facilitate the entrance of MMA in cancer cells, we depleted the old sera of lipids or of molecules larger than 3 kDa, two manipulations that should not affect the levels of polar metabolites such as MMA. In both cases, the ability of the depleted old sera to induce pro-aggressive properties was abolished (Extended Data Fig. 6e). Strikingly, both manipulations also caused a pronounced decrease in serum MMA levels (Extended Data Fig. 6f), indicating that the MMA responsible for this phenotype is complexed with a lipid moiety of a large size. These observations suggested to us that MMA may be complexed with lipidic structures (LSs) larger than 3 KDa in the sera thereby facilitating its entry into cancer cells. To test this hypothesis, we first complexed MMA with synthetic LSs or with LSs purified from fetal bovine serum (FBS). With both approaches, the concentration of MMA necessary to induce pro-aggressive properties was reduced to the levels similar to the old donor sera (Fig. 2h, Extended Data Fig. 6g, h). Moreover, MMA complexed with LSs from FBS produced a similar intracellular concentration of MMA within the same time frame as treatment with old donor sera (Fig. 2g). In support of this idea, treatment of cancer cells with LSs isolated from old sera, but not from young sera, or isolated from young sera and loaded with MMA at concentrations similar to the ones found in the old sera was sufficient to drive pro-aggressive properties (Fig. 2i). Conversely, LS depletion from old sera resulted in a reduction of total serum MMA levels and was sufficient to abrogate the pro-aggressive phenotype (Fig. 2j, k). Orthotopic injections of MDA-MB-231 cells into the mammary fat pads of athymic mice with elevated circulatory MMA levels further demonstrated a significant role of MMA in tumor progression by promoting tumor growth, metastatic spread and a concomitant significant decrease in survival of this cancer model (Fig. 2l-o, Extended Data Fig.5 j-l). Altogether, our data show that MMA, complexed with LSs, is a circulatory factor that contributes to the pro-aggressive effects of aging in cancer cells and is sufficient to drive tumor progression and aggressiveness.

To investigate how MMA promotes the cellular plasticity required to promote tumor progression, we performed a global transcriptomic analysis in NSCLC cells treated with MMA for 10 days. We found a dramatic transcriptional reprogramming induced by MMA (Fig. 3a, Extended Data Table 2). Functional annotation clustering analysis showed that MMA regulates genetic programs associated with cell fate decisions, such as wound healing and pattern specification (Fig. 3b), as well as genes involved in resistance to chemotherapeutic drugs, including several members of the ABC transporter family (Fig. 3c). Many of the upregulated genes encode secreted proteins known to remodel the tumor microenvironment, including factors that promote reorganization of the extracellular matrix, such as SERPINE1, MMP2 and CTGF^14,15,33^, immunosuppressive cytokines such as CXCL8, CCL2 and IL32^34,35^, as well as ligands that promote cell-to-cell communication such as WNT3, WNT5B and BMP8B^36,37^ (Fig. 3c, Extended Data Fig. 7a). Altogether, MMA appears to control a panoply of genetic programs, remodeling both the tumor and the microenvironment to promote aggressiveness and cancer progression.

**Fig. 3:**
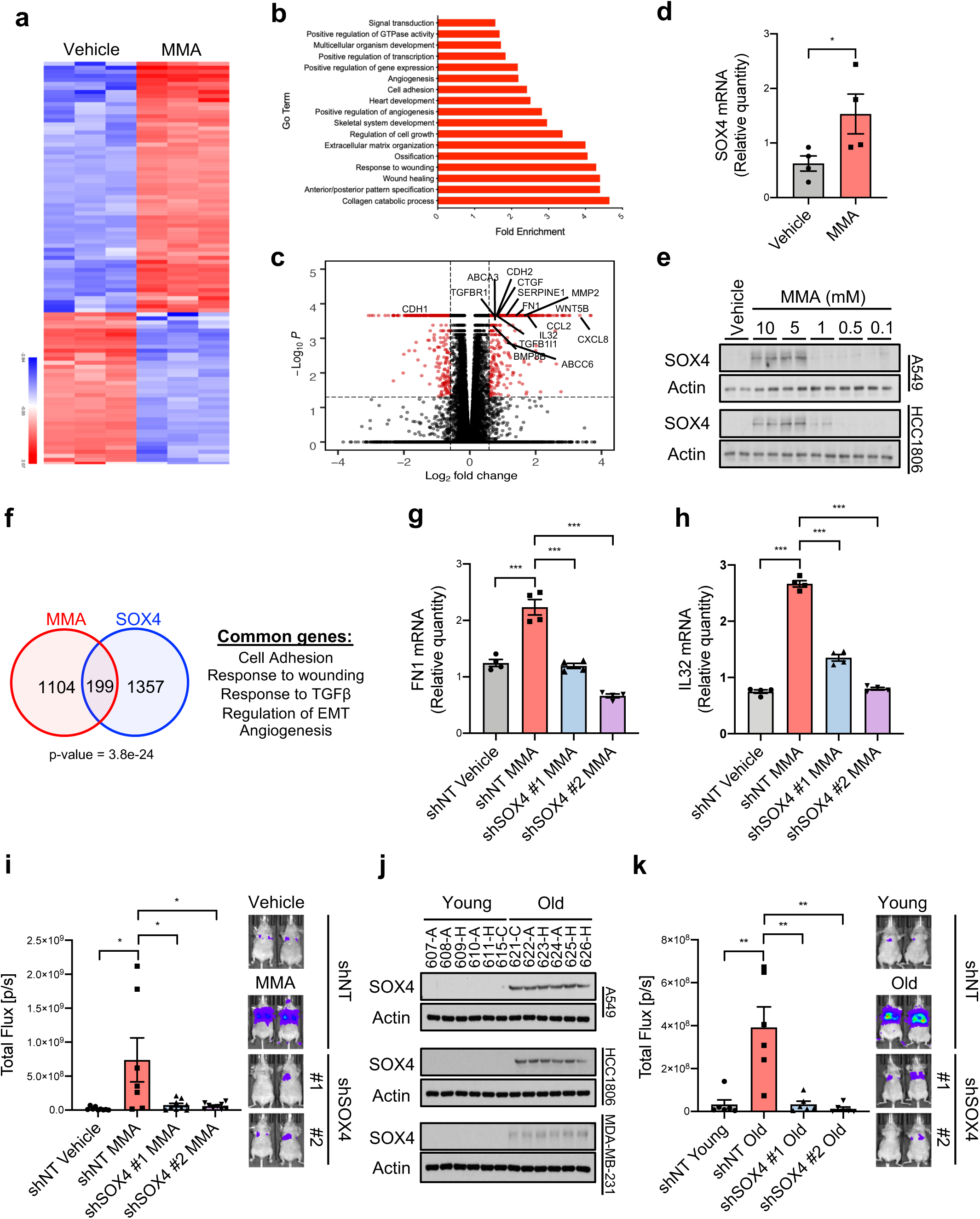
Methylmalonic acid triggers a global transcriptional reprogramming that underlies the acquisition of aggressive properties through induction of SOX4. **a-c**, Summary of RNA-seq analysis in A549 cells treated with 5 mM MMA for 10 days: a heatmap representation of hierarchical clustering of the top 100 changed mRNAs (**a**), functional annotation clustering analysis of the >1.5-fold changed mRNAs (**b**), and a volcano plot representation of the complete curated data set (the statistically significantly (FDR≤0.05) altered mRNAs that are changed more than 1.5-fold are displayed in red) (**c**) (n=3). **d**, mRNA levels of SOX4 evaluated by qPCR in A549 cells treated with 5 mM MMA for 3 days (n=4). **e**, Immunoblots for SOX4 in A549 and HCC1806 cells treated with MMA for 10 days; representative images (n=4). **f**, Venn diagram showing the overlap of altered mRNAs between A549 cells treated with MMA for 10 days and the altered genes induced by SOX4 induction^43^. **g, h**, mRNA levels of fibronectin (**g**) and IL32 (**h**) evaluated by qPCR in A549 cells with SOX4 knockdown and treated with 5 mM MMA for 10 days (n=4). **i**, Lung colonization assay of MDA-MB-231-luciferase cells with SOX4 knockdown and treated with 5 mM MMA for 5 days (n=8). **j**, Immunoblots for SOX4 in A549 cells cultured for 4 days in 10% human serum; A – African American; C – Caucasian; H – Hispanic (n=6). **k**, Lung colonization assay of MDA-MB-231-luciferase cells with SOX4 knockdown and treated with 10% human serum from old donors for 5 days (n=6). All values are expressed as mean ± SEM (*p<0.05, **p<0.01, ***p<0.001).

Gene annotation enrichment analysis also showed MMA to be a positive regulator of transcription (Fig. 3b), suggesting that the observed pro-aggressive transcriptional reprogramming may be mediated through this role. To find the transcriptional regulators involved, we again performed global transcriptomic analysis at an earlier time point of MMA treatment (day 3). Surprisingly, out of 439 induced genes (upregulated ≥1.5 fold), only 12 were transcription factors, 9 of which were significantly changed with qPCR validation (Extended Data Fig. 7b; Extended Data Table 2). One of the most upregulated of these transcription factors was the SRY-related HMG box (SOX) 4 (Fig. 3d). Notably, SOX4 is a poor prognosis marker that contributes to tumor progression and metastasis formation, with aberrantly high expression in a wide variety of aggressive cancers^38-41^, and is known to be a master regulator of EMT^42^. The levels of SOX4 were considerably increased in a variety of cell models treated with different MMA concentrations (Fig. 3e, Extended Data Fig. 7c, d), as well as in cells cultured with aged sera (Fig. 3j), supporting the idea that SOX4 mediates the pro-aggressive phenotype observed. A comparison between genes induced by SOX4^43^ and the ones induced by MMA treatment revealed a statistically significant overlap of 199 genes (Fig. 3f, Extended Data Table 2). Functional annotation clustering analysis revealed the overlapping genes – such as FN1 (Fibronectin), CDH2 (N-Cadherin), MMP2, IL32 and TGFB1I1 – to be related to aggressiveness properties. To better understand the relationship between MMA and SOX4, we used shRNA to suppress SOX4 expression. Upon SOX4 suppression, treatment with MMA failed to upregulate the mRNA levels of several of these genes (Fig. 3g, h, Extended Data Fig. 7e, f). Moreover, suppression of SOX4 blocked the ability of MMA or old sera to induce EMT and aggressive markers such as MMP2, CTGF, SERPINE1 (Extended Data Fig. 7g, h, 8c). Finally, SOX4 depletion fully abrogated the ability of MMA to promote migratory and invasive properties (Extended Data Fig. 7i), resistance to chemotherapeutic drugs (Extended Data Fig. 8a, b), and the ability of MDA-MB-231 cells to form colonies in the lungs of athymic mice upon treatment with MMA or with old sera (Fig. 3i-k).

Having shown that MMA promotes a pro-aggressive transcriptional remodeling through SOX4 induction, we next sought to understand how MMA induces SOX4 levels. TCA-related metabolites are known for their ability to regulate transcription through regulation of histone methylation levels^44^. In the cancer cell models used in this study, however, treatment with MMA did not change total levels of major histone modifications, suggesting that MMA works through a different mechanism to regulate SOX4 induction (Extended Data Fig. 8d, e). Alternatively, the TGFβ pathway regulates SOX4 levels^45^, and further analysis of the RNA-seq data showed an increase of several components of the TGFβ signaling pathway, including the upregulation of TGFB2 ligand. We confirmed the increase in the mRNA of TGFB2 (Fig. 4a), which correlated with an increase over time in the abundance of TGFβ-2 in the media of cancer cells treated with MMA (Fig. 4b). Moreover, and supporting the physiological relevance of these findings, analysis of the tumor tissues from mice with elevated circulatory MMA levels showed a significant upregulation of TGFB2 ligand, a concomitant induction of the TGFβ signaling and upregulation of SOX4 (Fig. 4c, d). Time course analysis showed that MMA robustly induced TGFβ signaling within 24 hours, during which an increase in SOX4 is observed, before any of the pro-aggressive markers are detected (Fig. 4e), suggesting a link between activation of TGFβ signaling, SOX4 induction and the acquisition of pro-aggressive properties driven by MMA. To test if activation of TGFβ signaling is responsible for SOX4 upregulation, we concurrently treated cancer cells with MMA and a TGFβ receptor inhibitor or a pan TGFβ neutralizing antibody. Both inhibition of the TGFβ receptor or neutralization of TGFβ ligand in the media was sufficient to block the ability of MMA to induce SOX4 and pro-aggressive properties (Fig. 4f, g). These data suggest that MMA relies on the activation of TGFβ signaling in an autocrine fashion to induce SOX4 and consequently the transcriptional reprogramming necessary for the cellular plasticity that sustains the metastatic cascade and tumor progression.

**Fig. 4:**
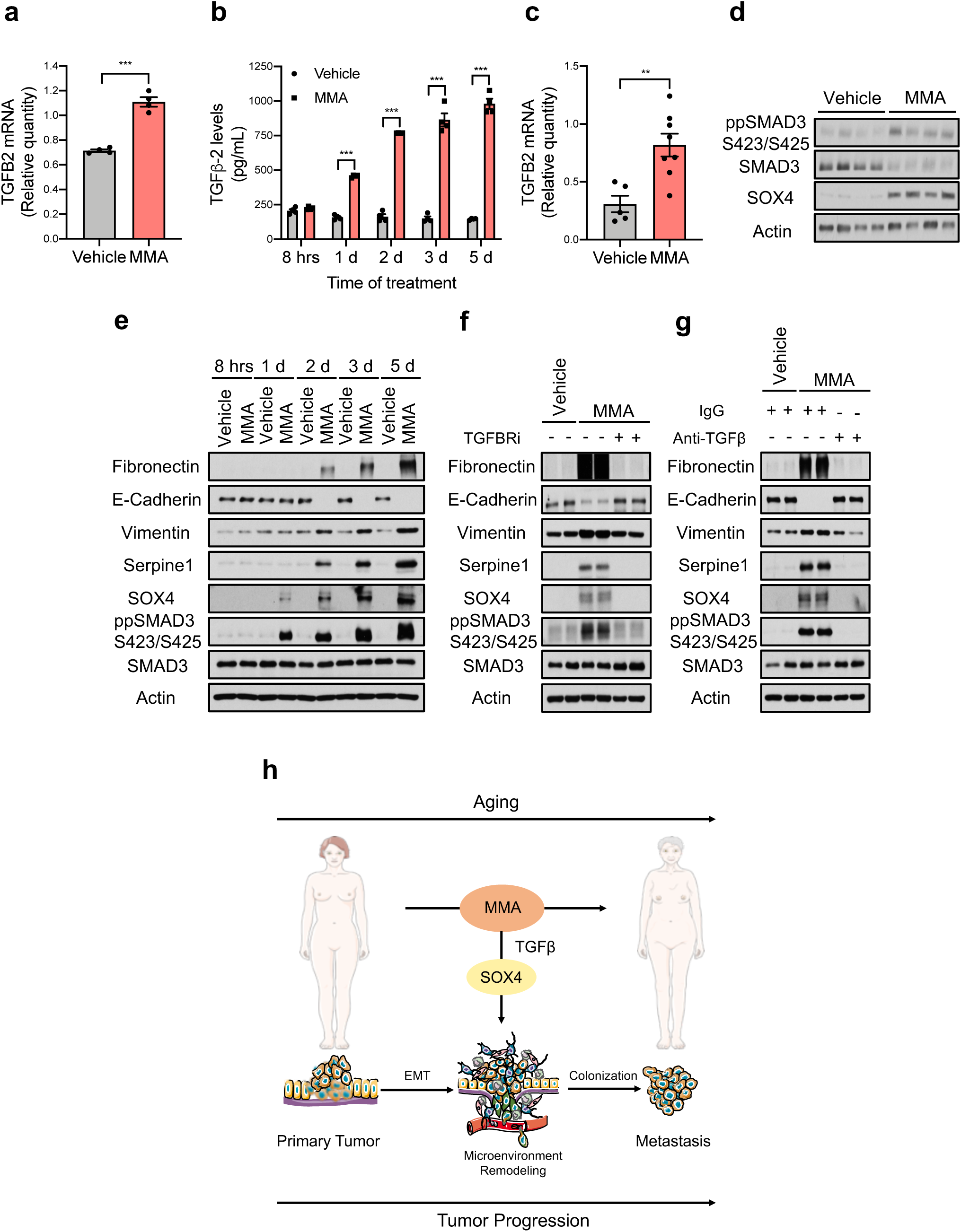
Methylmalonic acid induces SOX4 through activation of TGFβ signaling. **a**, TGFB2 mRNA levels determined by qPCR in A549 cells treated with 5 mM MMA for 3 days (n=4). **b**, Levels of TGFβ-2 ligand in conditioned media from A549 cells treated with 5mM MMA (n=4). **c, d**, TGFB2 mRNA levels determined by qPCR (vehicle n=5, MMA n=8) (c) and immunoblots for TGFβ pathway effectors and SOX4 (representative images, n=8) (d) in tumor samples from mice subcutaneously injected with the lower dose of MMA daily. **e-g**, Immunoblots for aggressiveness markers in A549 cells treated with 5 mM MMA (**e**), with 5 mM MMA in the presence of TGFBR inhibitor (**f**), or with 5 mM MMA in the presence of TGFβ neutralizing antibody (**g**); representative images (n=4). **h**, Age-induced accumulation of circulatory MMA induces SOX4 expression through the TGFβ pathway and elicits a transcriptional reprogramming that supports aggressiveness, promoting tumor progression and metastasis formation. All values are expressed as mean ± SEM (***p<0.001).

Taken together, our results show that metabolic deregulation of the aged host plays a central role in the acquisition of aggressive properties thus contributing to tumor progression. Specifically, aging promotes the increase in circulatory MMA, which in turn endows cancer cells with the properties necessary to migrate, invade, survive and thrive as metastatic lesions resulting in decreased cancer-associated survival (Fig. 4h). Although more in-depth studies are necessary to fully determine the scope of age-driven changes that contribute to the tumorigenic process, this study adds metabolic reprogramming to the complex relationship between aging and cancer.

## Contributions

A.P.G and J.B. conceived the project. A.P.G. and D.I. performed all the molecular biology experiments, the EMT-related experiments, the invasion and migration experiments, prepared the RNA for RNA-seq experiments and assisted on all other experiments. A.P.M. quantified the migration and invasion experiments. T.S. evaluated the stemness markers. V.L., J.E.E. and D.B. performed the mouse experiments. A.R. performed the drug resistance assays, prepared the metabolites for metabolomic analysis, produced the viral particles and assisted in multiple other experiments. J.H. and C.K. generated the constructs and the cell lines. C.K. and J.E.E. performed the histone post-translation modifications assessment. D.B and M.P. measured the concentration of MMA and analyzed the data. A.A and N.D. performed and analyzed the proteomic analysis of the human serum samples. T.S, E.M., I.E. and J.F.G assisted with the metabolomics experiments and helped with the metabolite treatments. J.A. performed the metabolomic analysis in the human sera. A.P.G., J.A., L.C.C., R.deC., N.D., S.M.F. and J.B. supervised the project. A.P.G., D.I., J.F.G., T.S., V.L., J.E.E., and N.D. analyzed the data. The manuscript was written by A.P.G. and edited by J.B., D.I., V.L., T.S., J.F.G., I.E., D.B. and S.M.F. All authors discussed the results and approved the manuscript.

## Acknowledgments

We are grateful to members of the Blenis and Cantley Laboratories for critical input on this project. We are also thankful to Dr. Paul Coffer for kindly providing the list of SOX4 targets and Robert Pritchard for experimental assistance. A.P.G. is supported by a Susan G. Komen Postdoctoral Fellowship and a Pathway to Independence Award from NCI (K99CA218686-01). T.S. is supported by the NIH F31 pre-doctoral fellowship 1F31CA220750-01. J.F.G. is supported by an FWO fellowship. C.K. was supported by a Medical Scientist Training Program grant from the NIGM/NIH T32GM007739 to the Weill Cornell/Rockefeller/Sloan Kettering Tri-Institutional MD-PhD Program. This research was supported by the NIH grant R01CA46595 to J.B. S.M.F. is funded by the European Research Council under the ERC Consolidator Grant Agreement number 711486 – MetaRegulation, FWO research grants and projects, and KU Leuven Methusalem Co-funding. L.C.C. owns equity in, receives compensation from, and serves on the Board of Directors and Scientific Advisory Board of Agios Pharmaceuticals and Petra Pharma Corporation. No potential conflicts of interest were disclosed by the other authors.

