## Extended Data Figure 4 for "Age-induced methylmalonic acid accumulation promotes tumor progression and aggressiveness"

**a**

**b**

**c**

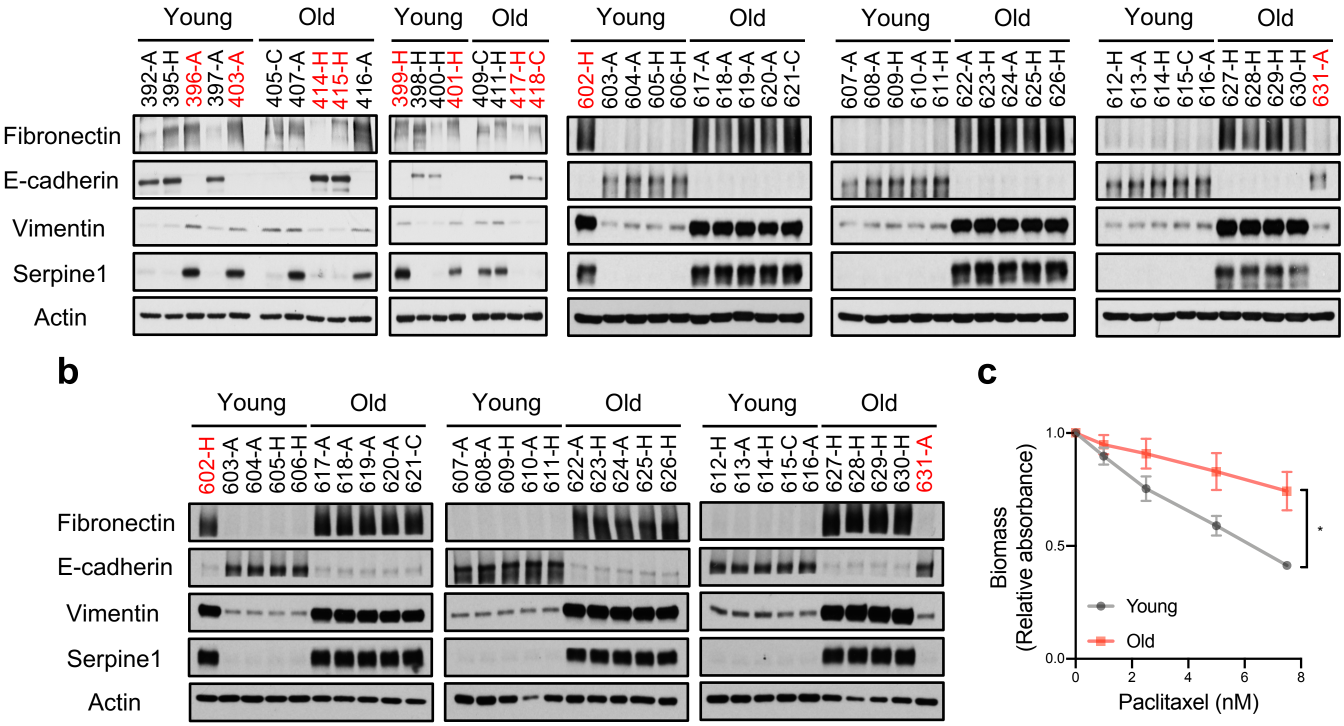


**d**

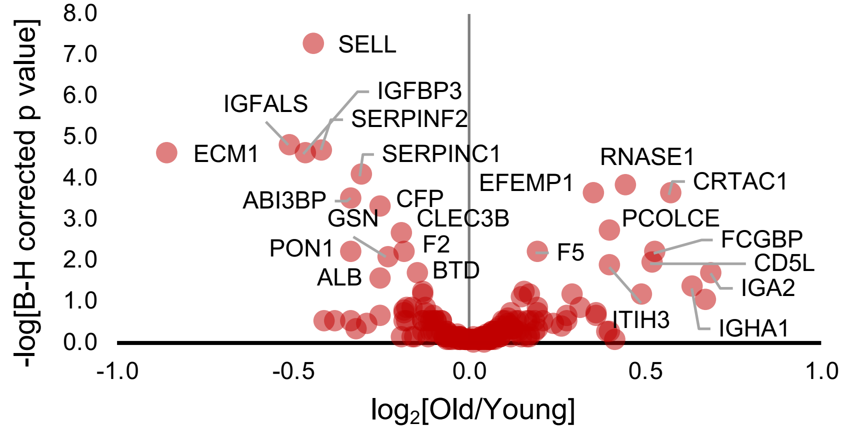


**e**

| Metabolite | Log2(FC) | -Log10(p) |
|---|---|---|
| Spermidine | -1.1827 | 1.1927 |
| Glutamine | -1.2089 | 1.5216 |
| Phenylpropiolic acid | -1.2597 | 1.5216 |
| α-Ketoglutarate | -1.2772 | 1.5216 |
| Glutathione disulfide | -1.312 | 2.9128 |
| 1,3-Diphosphoglycerate | -1.6237 | 1.5887 |
| Glutathione | -1.6851 | 3.6974 |

**f**

**g**

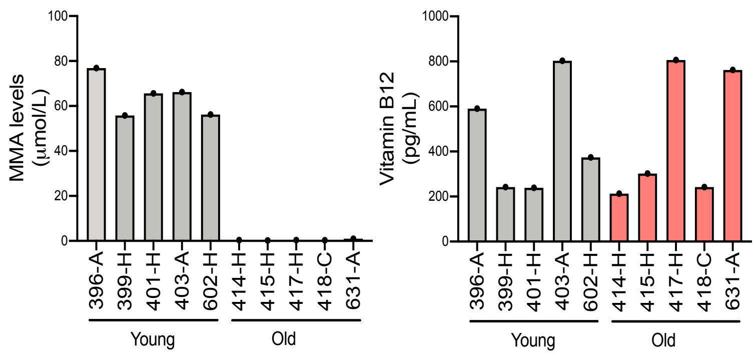
