## Extended Data Figure Legends for "Age-induced methylmalonic acid accumulation promotes tumor progression and aggressiveness"

**Extended Data Fig. 1 Serum of old donors induces a mesenchymal-like phenotype in non-small cell lung cancer cells.**  Morphology of A549 cells cultured for 4 days in 10% human serum; representative images of the first batch of 30 donors (n=15). A – African American; C – Caucasian; H – Hispanic; red label indicates outlier donors.

**Extended Data Fig. 2 Serum of old donors induces a mesenchymal-like phenotype in non-small cell lung cancer cells.**  Morphology of A549 cells cultured for 4 days in 10% human serum; representative images of the second batch of 30 donors (n=15). A – African American; C – Caucasian; H – Hispanic; red label indicates outlier donors.

**Extended Data Fig. 3 Serum of old donors induces a mesenchymal-like phenotype in triple negative breast cancer cells.**  Morphology of HCC1806 cells cultured for 4 days in 10% human serum; representative images of the second batch of 30 donors (n=15). A – African American; C – Caucasian; H – Hispanic; red label indicates outlier donors.

**Extended Data Fig. 4 Serum of the elderly induces** **aggressive properties in cancer cells and displays a distinct metabolic profile. a, b,** Immunoblots for aggressiveness markers in A549 (n=30) (**a**) and HCC1806 (n=15) (**b**) cells cultured for 4 days in 10% human serum; representative images **c,** Resistance to paclitaxel in A549 cells cultured for 4 days in 10% human serum (n=15). **d,** Volcano plot summarizing the proteomics analysis of all 60 human serum samples used in this study. **e,** List of all metabolites that are decreased at a statistically significant level in the sera of old donors (n=11). **f, g,** Concentrations of MMA (**f**) and vitamin B12 (**g**) in 10 outlier human sera (serum from 5 young and 5 old donors). For (**a, b, e, f):** A – African American; C – Caucasian; H – Hispanic. All values are expressed as mean ± SEM (*p<0.05).

**Extended Data Fig. 5 Methylmalonic acid promotes metastatic-like properties. a, b,** Immunoblots for aggressiveness markers in MCF-10A (**a**) and HCC1806 (**b**) cells treated with MMA for 10 days (n=4); representative images. **c,** Transwell migration assay of A549 cells treated with 5 mM MMA for 10 days (n=4). **d, e,** Stemness evaluated by the increase in the CD44 marker in MCF-10A cells treated with MMA for 10 days (**d**) and by the increase in the CD44 marker and the decrease in CD24 marker in A549 cells treated with 5 mM MMA for 10 days (**e**) (n=4). **f-h,** Resistance to paclitaxel in A549 (**f**) and HCC1806 cells (**g**) or to carboplatin in HCC1806 (**h**) cells treated with 5 mM MMA for 10 days (n=4). **i,** Immunoblots for aggressiveness markers in MDA-MB-231-luciferase cells treated with 5 mM MMA for 5 days; representative images (n=4). **j-l** End-point serum MMA concentrations (n=8) (**j**), bioluminescence intensity of the primary tumors (n=10) (**k**), and metastases (n=10) (**l**) in mice that were xenografted with MDA-MB-231-luciferase cells and treated with MMA in their drinking water. All values are expressed as mean ± SEM (**p<0.01, ***p<0.001).

**Extended Data Fig. 6 Methylmalonic acid delivery is regulated by lipidic structures (LSs) in the sera of old donors**

**a-d,** Immunoblots for aggressiveness markers in MCF-10A (**a, c**), A549 and HCC1806 (**b, d**) cells treated with 5 mM of the indicated acids (**a, b**) or with various concentrations (5–0.001 mM) of dimethyl methylmalonic acid (**c, d**) for 10 days; representative images (n=4). **e,** Immunoblots for aggressiveness markers in A549 cells cultured for 4 days with 10% young and old untreated human serum, or old serum that was passed through size exclusion columns or delipidated; representative images (n=6). **f,** MMA concentrations in the human serum used in (**e**) (n=6). **g, h,** Immunoblots for aggressiveness markers in A549 cells treated with complexes of lipofectamine (LA) with indicated amounts of MMA (n=3) (**g**) or in MCF-10A cells treated with LSs isolated from FBS that were complexed with indicated amounts of MMA (n=4) (**h**); representative images. All values are expressed as mean ± SEM (***p<0.001).

**Extended Data Fig. 7 Methylmalonic acid induces mRNA of pro-aggressive and poor prognosis genes. a.** mRNA levels of pro-aggressive cell intrinsic factors and secreted factors evaluated by qPCR in A549 cells treated with 5 mM MMA for 10 days (n=4). **b,** mRNA levels of transcription factors evaluated by qPCR in A549 cells treated with 5 mM MMA for 3 days (n=3). **c, d,** Immunoblots for SOX4 in MCF-10A cells treated with MMA for 10 days (**c**) and in MDA-MB-231-luciferase cells treated with 5 mM MMA for 5 days (**d**); representative images (n=4). **e, f,** mRNA levels of N-cadherin (CDH2) (**e**) and TGFB1I1 (**f**) evaluated by qPCR in A549 cells with SOX4 knockdown and treated with 5 mM MMA for 10 days (n=4). **g, h,** Immunoblots for aggressiveness markers in A549 (**g**) and MDA-MB-231-luciferase cells (**h**) with SOX4 knockdown and treated with 5 mM MMA for 10 days and 5 days respectively; representative images (n=4). **i,** Transwell migration/invasion assays of MDA-MB-231-luciferase cells with SOX4 knockdown and treated with 5 mM MMA for 5 days (n=4). All values are expressed as mean ± SEM (*p<0.05, **p<0.01, ***p<0.001).

**Extended Data Fig. 8 Methylmalonic acid does not alter levels of major histone post-translational modifications**

**a, b,** Resistance to carboplatin and paclitaxel in A549 cells with SOX4 knockdown and treated with 5 mM MMA for 10 days (**a**) and in MDA-MB-231-luciferase cells with SOX4 knockdown and treated with 5 mM MMA for 5 days (**b**) (n=4). **c,** Immunoblots for aggressiveness markers in MDA-MB-231-luciferase cells with SOX4 knockdown and treated with sera from young and old donors for 5 days; representative images (n=6) **d, e,** Immunoblots for histone marks in A549 (**c**) and MCF-10A (**d**) cells treated with 5 mM MMA for 1 or 3 days; representative images (n=4). All values are expressed as mean ± SEM (*p<0.05, **p<0.01, ***p<0.001).
