## Supplementary material for "Age-induced methylmalonic acid accumulation promotes tumor progression and aggressiveness": Materials and Methods

**Online Methods**

**Cell Lines**

MCF-10A human mammary epithelial cells were obtained from the American Type Culture Collection (ATCC), and were cultured as previously described^1^. HCC1806 human breast cancer and A549 human lung cancer cell lines were also obtained from ATCC and were cultured in RPMI 1640 medium (Corning) supplemented with 10% FBS (Sigma-Aldrich) and penicillin-streptomycin (Gibco). MDA-MB-231-luciferase cells described previously^2^ were obtained from Dr. Massague’s lab and were maintained in high glucose DMEM (Gibco) supplemented with 10% FBS (Sigma-Aldrich) and penicillin-streptomycin (Gibco). HEK293T cells were obtained from GenHunter and cultured in high glucose DMEM (Gibco) supplemented with 10% FBS (Sigma-Aldrich) and penicillin-streptomycin (Gibco). All cell lines were maintained at 37°C and 5% CO_2_. All cell lines were routinely tested for mycoplasma and were at all times mycoplasma negative.

### **Mice**

Female nu/nu athymic mice were purchased from Envigo at the age of 4-6 weeks. Xenograft experiments were started 7-10 days after the mice were received. The mice were maintained at Weill Cornell Medicine in compliance to Weill Cornell Medicine Institutional Animal Care and Use Committee protocols.

**Human Serum**

Human serum from 30 young (aged 30 and below) and 30 old (aged 60 and above) male individuals with no diagnosed disease at the time of collection were obtained from BioreclamationIVT collected as two separate batches (15 young and 15 old donors in each batch)**.** The specific serum used in this study is limited but samples from similar donors can be obtained from BioreclamationIVT. For detailed information on the donors please see Extended Data Table 1 for details.

**METHOD DETAILS**

**Cell Culture Treatments**

To test the effects of aged serum in cancer cells, A549 or HCC1806 cells were seeded and their media replaced the next day with media containing 10% human serum for 4 days. Prior to replacement of the culturing media, the cells were washed 3 times with PBS. To evaluate the effects of the top upregulated metabolites in the serum of the elderly, A549 cells were plated in normal culture media and the following day treated with 5 mM quinolinate (Sigma-Aldrich), 5 mM phosphoenolpyruvate (Sigma-Aldrich), 5 mM methylmalonic acid (MMA; Tocris), or vehicle (0.1% DMSO for quinolinate; double distilled water for MMA and phosphoenolpyruvate). For the subsequent treatments to determine the effects of MMA on cellular phenotypes, MCF-10A, HCC1806, MDA-MB-231-luciferase and A549 cells were seeded and the next day treatment was initiated with indicated concentrations of MMA (Tocris) or vehicle (double distilled water) for the time frames indicated. To test the specificity of MMA treatments in the phenotypes tested, A549 and HCC1806 cells were treated with 5 mM MMA (Tocris), 5 mM malonic acid (Sigma-Aldrich), 5 mM propionic acid (Sigma-Aldrich), 5 mM fumaric acid (Sigma-Aldrich), 5 mM pyruvic acid (Sigma-Aldrich), or vehicle (0.1% DMSO) for 10 days. MCF-10A cells were treated with vehicle (0.1% DMSO) or 1 mM each of the following acids: MMA (Tocris), malonic acid, propionic acid, succinic acid, pyruvic acid, hydroxyisobutyric acid (Sigma-Aldrich), fumaric acid (Sigma-Aldrich), maleic acid (Sigma-Aldrich), malic acid (Sigma-Aldrich), or ⍺-ketoglutaric acid (Sigma-Aldrich) for 10 days. To test the effects of enhanced MMA permeability A549 and HCC1806 cells were treated with 5 mM MMA or a range of dimethyl-MMA (Sigma-Aldrich) concentrations (5, 0.5, 0.05 or 0.005 mM), or vehicle (0.1% DMSO) for 10 days. MCF-10A cells, on the other hand, were treated with 1 mM MMA or a range of dimethyl-MMA concentrations (1, 0.1, 0.01 or 0.001 mM), or vehicle (0.1% DMSO) for 10 days. To test the role of TGFβ signaling in SOX4 induction upon MMA treatment, A549 cells were pretreated for 2 hours with TGFβ receptor inhibitor SB431542 (S1067; Selleck Chem) dissolved in DMSO or with 0.5 µg/ml TGFβ neutralizing antibody (MAB1835, R&D Biosystems; normal mouse IgG from Santa Cruz sc2025 used as control), and then maintained with the inhibitor or the antibody for the duration of MMA treatment. For all acidic treatments, 25 mM HEPES (Sigma-Aldrich) was added to the treatment media to buffer potential changes in pH, and the media were replaced every day during the treatments.

**Targeted Metabolomics and Data Analysis**

Circulatory polar metabolites were extracted using 80% (v/v) aqueous methanol as described before^3^ from 100 μL of sera of young and old donors. Targeted liquid chromatography-tandem mass spectrometry (LC-MS/MS) was performed using a 5500 QTRAP triple quadrupole mass spectrometer (AB/SCIEX) coupled to a Prominence UFLC HPLC system (Shimadzu) with Amide HILIC chromatography (Waters). Data were acquired in selected reaction monitoring (SRM) mode using positive/negative ion polarity switching for steady-state polar profiling of greater than 260 molecules. Peak areas from the total ion current for each metabolite SRM transition were integrated using MultiQuant v2.0 software (AB/SCIEX). Statistical analysis of the data was carried out using MetaboAnalyst, a free online software for the analysis of metabolomic experiments ([www.metaboanalyst.ca](http://www.metaboanalyst.ca)). The original data was normalized to the median of the entire metabolome in each sample and log transformed prior to further analysis (Extended Data Table 1).

**Proteomic Analysis of Human Serum**

Abundant serum proteins were depleted using High Select Top 14 spin columns (Thermo #A36370) following the manufacturer provided protocol. Briefly, 10 µl of serum was applied to each column and incubated for 10 min with end-over-end rotation. Depleted samples were collected by centrifugation at 2000 x g for 2 min. Ice-cold 100% trichloroacetic acid was added to 20% final concentration. Proteins were allowed to precipitate on ice for 60 min and then pelleted for 10 min at 20k x g at 4°C. Pellets were washed twice with ice-cold acetone and allowed to dry at room temperature. Dry protein pellets were re-suspended in 8 M urea, 50 mM ammonium bicarbonate (ambic). Proteins were reduced by addition of dithiothreitol (DTT) to 5 mM and incubating at room temperature for 30 min, then alkylated by adding iodoacetamide to 15 mM and incubating in the dark at room temperature for 30 min. Iodoacetamide was quenched with an additional 5 mM DTT. Samples were diluted to 2 M urea with 50 mM ambic and digested overnight at room temperature by adding 600 ng Lysyl endopeptidase (lysC, Wako Chemicals USA, Inc.). Samples were further diluted to 1 M urea with 50 mM ambic and digested with 600 ng sequencing grade modified trypsin (Promega) for 6 hr at 37°C with shaking. Digests were acidified by the addition of neat formic acid (FA) to 2% final concentration, and desalted on hand packed C18 STAGE Tips^4^. Eluted peptides were dried in a centrifugal evaporator. Peptides were labeled with 10-plex amine reactive TMT labeling reagents (Thermo Fisher, Rockford, IL) by re-suspending in 100 µl of 0.2 M HEPES pH 8 and adding 0.2 mg of each TMT label in 10 µl of anhydrous acetonitrile and incubating at RT for 1 hr. Reactions were quenched with 8 µl of 5% hydroxylamine and then acidified with 16 µl neat FA. Test mixtures for each 10plex set were generated by mixing 5 µl from each channel and analyzed with a 75 min gradient version of the final analysis method (described below). The final mix was adjusted based on this analysis to generate equal total reporter ion intensities from each TMT channel. Mixed peptides were desalted on 50 mg tC18 Sep-Pak cartridges (Waters, Milford, MA), dried, and re-suspended in 5 µl of 5% FA. Mass Spectrometric analysis was performed on a Thermo Orbitrap Fusion mass spectrometer (Thermo Fisher, Waltham, MA) equipped with an Easy nLC-1000 UHPLC (Thermo Fisher Scientific). Peptides were separated with a gradient of 6–25% ACN in 0.1 % FA over 155 min and introduced into the mass spectrometer by nano-electrospray as they eluted off a self-packed 40 cm, 75 µm (ID) reverse-phase column packed with 1.8 µm, 120 Å pore size, C18 resin (Sepax Technologies, Newark, DE). They were detected using a data-dependent MS2 method with a real-time search (RTS) plugin^5^ used to trigger MS3 scans for TMT reporter ion quantification. For each cycle, one full MS scan was acquired in the Orbitrap at a resolution of 120,000 with automatic gain control (AGC) target of 5 × 10^5^ and a maximum ion accumulation time of 100 ms. Each full scan was followed by the selection of the most intense ions, as many as possible in 2 s total cycle time, for collision induced dissociation (CID) and MS2 analysis in the linear ion trap for peptide identification using an AGC target of 1.5 × 10^4^ and a maximum ion accumulation time of 50 ms. Ions selected for MS2 analysis were excluded from reanalysis for 60 s. Ions with +1 or unassigned charge were also excluded from analysis. MS2 spectra were searched in real-time using the RTS module^5^. RTS settings required a binomial score threshold of 65 to trigger SPS MS3 scans using positively identified MS2 fragment ions. Selected MS2 ions were fragmented with a HCD collision energy of 55 and scanned in the orbitrap at a resolution of 50,000 at m/z 200. To increase coverage of lower abundance proteins, we used the gene close-out feature to trigger a maximum of 10 MS3 per protein. MS/MS spectra were matched to peptide sequences using SEQUEST v.28 (rev. 13)^6^ and a composite database containing the 20,415 Uniprot reviewed canonical predicted human protein sequences (<http://uniprot.org>, downloaded 5/1/2019) and its reversed complement. Search parameters allowed for three missed cleavages, a mass tolerance of 20 ppm, a static modification of 57.02146 Da (carboxyamidomethylation) on cysteine, and dynamic modifications of 15.99491 Da (oxidation) on methionine and 229.16293 for TMT on lysines and peptide amino termini. Peptide spectral matches (PSMs) were filtered to 1% FDR using the target-decoy strategy^7^ combined with linear discriminant analysis (LDA)^8^ using the SEQUEST Xcorr and ΔCn' scores, precursor mass error, observed ion charge state, and the number of missed cleavages. The data were further filtered to a 1% protein FDR using the same strategy with protein scores derived from the product of all LDA peptide probabilities. Remaining peptide matches to the decoy database as well as contaminating proteins (e.g., human keratins) were removed from the final data set. TMT reporter ion signal-to-noise (SN) values were extracted for all PSMs by identifying the maximum peak intensity within a 3 millidalton window around the theoretical m/z. Each PSM was required to have a sum reporter ion SN across all 10 TMT channels ≥ 100 for inclusion in subsequent protein quantification. Reporter ion intensities were adjusted to correct for the isotopic impurities of the different TMT reagents based on manufacturer supplied values. Protein quantification was performed separately for each 10plex by summing SN values from all matching PSMs for each channel. The protein sum SN values were normalized to correct for mixing errors by dividing each value by the sum of all values within its channel. These values were then transformed for each protein to generate a fractional intensity for each sample. All raw data files, peak lists, and the sequence database have been deposited in the MASSive repository (<https://massive.ucsd.edu>, ID#: MSV000084974) and are available for download at ftp://massive.ucsd.edu/MSV000084974.

**Measurements of MMA and Vitamin B12 Concentrations in Human Serum**

Frozen aliquots of the human serum (unprocessed, delipidated, size excluded or lipidic structure depleted) were sent to ARUP Laboratories (Sat Lake City, Utah) for measurement of MMA (test code: 2005255) and vitamin B12 (test code: 0070150) concentrations. ARUP Laboratories is a national nonprofit and academic reference laboratory of diagnostic medicine.

**Delipidation and Size-Exclusion in Human Serum**

Human serum samples were manipulated to assess the components of human serum that might facilitate entrance of MMA into cells. To delipidate the human serum Cleanascite Lipid Removal Reagent (Biotech Support Group) was utilized according to the manufacturer’s protocol specifically for “serum” samples, using a 1:4 volume ratio of Cleanascite reagent to sample. To deplete the serum from molecules larger than 3 kDa serial filtration through size exclusion columns was performed. Initially sera was applied to Amicon Ultra-4 centrifugal filter units (Milipore) with a molecular size cut-off of 100 kDa and centrifuged at 4000 x g at 4°C. The flow-through fraction then was processed successively through filter units with molecular size cut-offs of 50, 10 and 3 kDa. Delipidated or size-excluded human serum fractions were used in cell culture treatments similarly to unprocessed human serum as described above.

**Lipidic Structure (LS) Isolation from Serum and LS Depletion from Human Serum**

Lipidic structures were isolated from freshly thawed fetal bovine serum (FBS) or human serum using Total Exosome Isolation (from Serum) reagent (Invitrogen) according to the manufacturer’s protocol. Briefly 6 ml FBS or 1.5 ml of each human serum was pelleted with the reagent (supernatants were saved to be used as LS-depleted serum) and then the pellets were resuspended in sterile PBS in a volume equal to half of the starting serum volume (3 ml and 750 ul respectively). Aliquots of LSs were kept at -80°C to prevent freeze thaw cycles. Cells were treated with 150 μl LSs in PBS from human serum (LSs from equivalent to 300 μl human serum) or 300 μl LS-depleted human serum in 6 cm plates with 3 ml normal growth media for a total of 4 day treatment. The media was changed and treatment repeated for a second time 48 hours after the initial treatment. To control for presence of the Total Exosome Isolation reagent in these LS-depleted sera, we added same amount of the reagent to cells treated with the control human serum.

**Complexing of MMA with Artificial or Serum-Derived Lipidic Structures (LSs)**

MMA was complexed with artificial LSs (Lipofectamine 2000) or with serum-derived LSs to achieve final concentrations of 1, 10 or 100 μM in the culture media. Briefly Lipofectamine 2000 and indicated amounts of MMA were diluted in Opti-MEM (Gibco) medium and incubated for half an hour for complex formation. A549 cells were treated with the complexes and the media were replaced with fresh growth media 24 hours later. Cells were treated one more time with freshly made complexes 48 hours after the initial treatment. The media were again replaced 24 hours later and cell lysates were collected 4 days after the initial treatment with the complexes. For complexing MMA with LSs isolated from either FBS or human serum Exo-Fect Exosome Transfection Kit (System Biosciences) was used according to the manufacturer’s protocol. Briefly 200 μl LSs from FBS or 300 μl LSs from each human serum was processed and resuspended in 600 μl PBS to be used to transfect 2-6 cm plates. Complexes were made fresh before the initial treatment and then the remaining complexes were stored at -80°C until they were used for the second treatment. Cells were treated with 300 μl MMA-complexed LSs in PBS in their normal growth media that contained either FBS or horse serum. The cells were treated with the remaining 300 μl MMA-complexed LSs a second time 48 hours after the initial treatment. The cell lysates were collected 4 days after the initial treatment.

**SOX4 Gene Silencing**

shSOX4 #1 (TRCN0000018213), shSOX4 #2 (TRCN0000018214) and shNT (shGFP - TRCN0000072181, all from Sigma Aldrich) lentiviruses were produced by co-transfection of HEK293T cells with plasmids encoding psPAX2 (Addgene plasmid 12260), and pMD2.G (Addgene plasmid 12259) using X-tremeGENE HP (Roche) in accordance with the manufacturer’s protocol. Medium was changed 24 hours post-transfection and the virus harvested after 48 hours, filtered, and used to infect MDA-MB-231 luciferase and A549 cells in the presence of 8 μg/mL polybrene (Sigma-Aldrich). Selection of resistant colonies was initiated 24 hours later using 2 μg/mL puromycin (Sigma-Aldrich) for 24 hours after which the cells were treated with MMA as described above for the time frame indicated.

**Quantification of TGFβ-2 Ligand Levels in Cell Culture Media**

A549 cells were treated with 5 mM MMA for indicated amounts of time and then the conditioned culture media were collected. The samples were centrifuged for 3 minutes at 300 g to remove any cells and debris and activated using the Sample Activation Kit 1 (R&D Systems) according to the manufacturer’s protocol. Human TGFβ-2 levels in these conditioned media were measured with the Human TGFB2 Quantikine ELISA Kit (R&D Systems) according to the manufacturer’s protocol.

**Chemotherapeutic Drug Assays**

A549, HCC1806 and MCF-10A cells were treated with the indicated concentrations of MMA or vehicle (double distilled water) for 10 days. A549 and MDA-MB-231-luciferase cells with SOX4 silenced were also treated with MMA as described above after which they were seeded in 96-well plates in technical triplicates. The cells were treated the next day with either vehicle control (DMSO (0.1%)), carboplatin (0-200 µM), or paclitaxel (0-7.5 nM) at various concentrations. The media containing the treatments were replaced every day for 4 days. At the end of the treatments the cells were fixed in 4% paraformaldehyde (Electron Microscopy Sciences) diluted in PBS for 30 minutes. After removing the fixative solution the plates were washed with PBS and stained with 0.1% Crystal Violet solution for 15 minutes. The staining solution was removed and the plates were washed 3 times under running water, to remove the excess stain, and allowed to dry at room temperature. To quantify the biomass, crystal violet staining was eluted with 100% methanol and the absorbance at 590 nm was measured using an Envision plate reader (Perkin Elmer).

**Analysis of CD24 and CD44**

Cells were dissociated using Cell Stripper (Corning), collected on ice and pelleted by centrifugation. After removing the Cell Stripper and washing the cell pellet with ice cold PBS, the cells were stained on ice for 30 minutes in 100 μL DMEM/F12 (without phenol red) with an APC mouse anti-human CD44 (559942, BD Biosciences) and FITC mouse anti-human CD24 (555427, BD Biosciences), or an APC mouse IgG2b (555745, BD Biosciences) and FITC mouse IgG2a (553456, BD Biosciences) as isotype controls. After labeling, each sample was washed twice with ice cold PBS and resolved on a BD Accuri B6 flow cytometer (BD Biosciences). Data analysis to determine the medium fluorescence intensity (MFI) of CD24 and CD44 positive cells was performed using the FlowJo software package. An example of the gating strategy performed prior to MFI analysis is shown below.

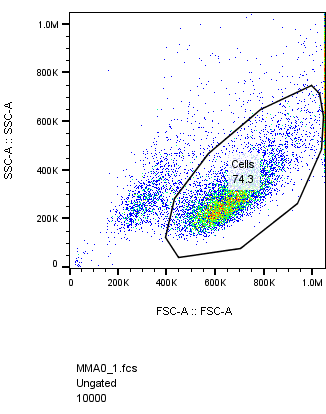

**MCF-10A**

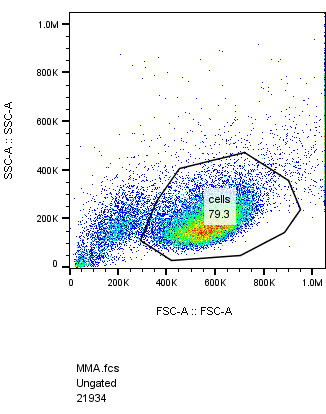

**A549**

**Transwell Migration and Invasion Assays**

MCF-10A cells were trypsinized and collected as previously described^9^. Resuspension media were aspirated and cells were resuspended in assay medium (DMEM/F12 (Corning), 0.5% Horse Serum (Gibco), 500 ng/ml hydrocortisone (Sigma), 100 ng/ml cholera toxin (Sigma)). For migration assays, Boyden chamber inserts (BD Biosciences, 8 μm pore size) were pre-coated with 25 µg/µl rat tail collagen 1 (Corning). Assay medium supplemented with 5 ng/ml EGF (Peprotech) was added to the bottom chamber of the cell culture inserts. Cells (5 x 10^4^ cells/ 250 μl assay media) were then added to the top chamber of cell culture inserts in a 24-well companion plate. After 6 hours of incubation, the cells that had migrated to the lower surface of the membrane were fixed with ethanol and stained with 0.2% crystal violet in 2% ethanol. For cell invasion assays, BD BioCoat invasion chambers coated with growth factor reduced Matrigel were used. Invasion chambers were prepared according to manufacturer’s specifications and assays were performed as described for migration assays except that 20 ng/ml of EGF (Peprotech) was added to MCF-10A assay media to serve as the chemo-attractant and cells were allowed to invade for 24 hours.

For MDA-MB-231 and A549, transwell migration and invasion assays were performed as described above with minor changes. For MDA-MB-231 cells, high-glucose DMEM (Gibco) supplemented with 250 μg/mL BSA (Sigma-Aldrich) was used as the assay medium, and high glucose DMEM media supplemented with 10% FBS (Sigma-Aldrich) was used as the chemoattractant for both migration (6 hours) and invasion assays (20 hours). For A549 cells, RPMI (Corning) supplemented with 250 μg/mL BSA (Sigma-Aldrich) was used as the assay medium, and RPMI media supplemented with 10% FBS (Sigma-Aldrich) was used as the chemoattractant for the 24-hour migration assay. Images of crystal violet stained cells were captured using a Nikon DS-Fi2 camera, and quantifications were carried out in an automated way using Fiji/ImageJ. Briefly, binary images of the area covered by crystal violet-positive cells was generated using thresholding and settings that were appropriate for control samples, and these settings were used throughout the analysis. The percentage area covered by crystal violet-positive cells was quantified for each condition, using a minimum of three technical replicates.

**Immunoblots for total cell lysates**

Proteins were isolated directly from intact cells via acid extraction using a 10% TCA solution (10% trichloroacetic acid, 25 mM NH_4_OAc, 1 mM EDTA, 10 mM Tris⋅HCl pH 8.0). Precipitated proteins were harvested and solubilized in a 0.1 M Tris⋅HCl pH 11 solution containing 3% SDS and boiled for 10-15 minutes. Protein content was determined with the DC Protein Assay kit II (BioRad), and 20 μg total protein from each sample were run on SDS-PAGE under reducing conditions. The separated proteins were electrophoretically transferred to a nitrocellulose membrane (GE Healthcare), which was blocked in TBS-based Odyssey Blocking buffer (LI-COR). Proteins of interest were probed with specific antibodies: E-cadherin (610181 - BD Biosciences), ZO1 (5406S - Cell Signaling), fibronectin (ab2413 - Abcam), vimentin (5741S - Cell Signaling), Serpine1 (612024 - BD Biosciences), CTGF (ab6992 - Abcam), MMP2 (4022S - Cell Signaling), SOX4 (ab8026 - Abcam), SMAD3 (9523S - Cell Signaling), ppSMAD3 S423/S425 (ab52903 - Abcam) and actin (sc1615 - Santa Cruz). Membranes were incubated with the primary antibodies overnight at 4°C, and then with the appropriate horseradish peroxidase–conjugated (HRP) anti-rabbit, anti-mouse (both from GE Healthcare), or anti-goat (Millipore) immunoglobulin for 2 hours at room temperature. The signals were developed using Amersham ECL detection system (GE Healthcare).

**Analysis of Histone Post-Translational Modifications**

A549 and MCF-10A cells were trypsinized and normalized for cell numbers. Cell pellets were washed twice with ice-cold PBS+ (PBS containing containing 5 mM sodium butyrate, 2 mM nicotinamide, 2 mM phenylmethylsulfonyl fluoride (PMSF), 10 mM Aprotinin, 10 mM Leupetin, 10 mM Pepstain A, 10 mM NaF, 10 mM NaVO_4_, 0.02% (w/v) NaN_3_) and then resuspended in Triton Extraction Buffer (TEB: PBS+ containing 0.5% Triton X 100 (v/v)) at a cell density of 10^7^ cells per ml. Cell lysis was achieved on a rotator at 4°C for 10 min. Nuclei were centrifuged at 6,500 x g for 10 min at 4°C, and washed once with half the volume of TEB. Histones were extracted in 0.2 N HCl at a density of 4x10^7^ nuclei per ml overnight at 4°C. To remove debris samples were centrifuged at 6,500 x g for 10 min at 4°C. The histone proteins present in the supernatant were collected and neutralized with 2 M NaOH at 1/10 of the volume of the supernatant. 8 µg of each sample was resolved on a 15% polyacrylamide gel and immunoblots were performed as described above. The following primary antibodies were utilized: H3K4me3 (61379 - Active Motif), H3K27me3 (39155 - Active Motif), H3K27ac (39133 - Active Motif), H4K8ac (ab15823 - Abcam), H3K56ac (39281 - Active Motif), H3K9ac (ab4441 - Abcam), H3K9me3 (ab8898 - Abcam), H3K36me3 (ab9050 - Abcam), Total H4 (ab10158 - Abcam), Total H3 (4499s - Cell Signaling).

**Gene Expression Analysis**

RNA was isolated using the PureLink RNA isolation kit (Life Technologies) and contaminant DNA was digested with DNAse I (Amplification grade, Sigma-Aldrich). cDNA was synthesized using the iSCRIPT cDNA synthesis kit (BioRad) and analyzed by quantitative PCR (qPCR) using SYBR green master mix (Life Technologies) on a QuantStudio6 Real-Time PCR system (Life Technologies). Target gene expression was normalized to Tata Binding Protein (*TBP*) and actin (*ACTB*) expression. Primer sequences can be found in the following table:

**Table:** Human Primers Used for qPCR Analysis, Related to Experimental Procedures

| **Gene** |  | **Primer Sequence** |
| --- | --- | --- |
| *TBP* | Forward | GAGCCAAGAGTGAAGAACAGTC |
|  | Reverse | GCTCCCCACCATATTCTGAATCT |
| *ACTB* | Forward | CATGTACGTTGCTATCCAGGC |
|  | Reverse | CTCCTTAATGTCACGCACGAT |
| *ABCA3* | Forward | CATCTCTACCATCTCCTTCAGC |
|  | Reverse | CAGAGTCATCCAGTTGTACCG |
| *CDH1* | Forward | CCCACCACGTACAAGGGTC |
|  | Reverse | CTGGGGTATTGGGGGCATC |
| *CDH2* | Forward | CCCAAGACAAAGAGACCCAG |
|  | Reverse | GCCACTGTGCTTACTGAATTG |
| *FN1* | Forward | CAGTGGGAGACCTCGAGAAG |
|  | Reverse | TCCCTCGGAACATCAGAAAC |
| *TGFBR1* | Forward | ACATGATTCAGCCACAGATACC |
|  | Reverse | GCATAGATGTCAGCACGTTTG |
| *WNT5B* | Forward | AAGGAGTTTGTGGATGCCC |
|  | Reverse | GCTACGTCTGCCATCTTATACAC |
| *TGFB1I1* | Forward | CTCTGTGAGCTAGATCGGTTG |
|  | Reverse | GGAGGCTGGGTCTTTTCTTATC |
| *TGFB2* | Forward | GAGGTTTACAAAATAGACATGCCG |
|  | Reverse | ACTCTGAACTCTGCTTTCACC |
| *MMP2* | Forward | ATGCCGCCTTTAACTGGAG |
|  | Reverse | GGAAGCCAGGATCCATTTT |
| *IL32* | Forward | CCTTGGCTCCTTGAACTTTTG |
|  | Reverse | CTGTCCACGTCCTGATTCTG |
| *CXCL8* | Forward | ACAAGCTTCTAGGACAAGAGC |
|  | Reverse | GCACTCCTTGGCAAAACTG |
| *CCL2* | Forward | CAGAAGTGGGTTCAGGATTCC |
|  | Reverse | ATTCTTGGGTTGTGGAGTGAG |
| *FOXL1* | Forward | CTTTCAACGCTTCCCTGATG |
|  | Reverse | CCGTGCCATTGTTTGCTTTA |
| *FOXS1* | Forward | GGACGCCAGGAATGTTCTT |
|  | Reverse | TGTGGTTGTCCTTGGCTC |
| *HOXD9* | Forward | GGCTGTTCGCTGAAGGAG |
|  | Reverse | CGTCTGGTATTTGGTGTAGGG |
| *HOXD10* | Forward | AGGTCTCCGTGTCCAGTC |
|  | Reverse | GCGTTTGGTGCTTAGTGTAAG |
| *JUNB* | Forward | GGACACGCCTTCTGAACG |
|  | Reverse | CGGAGTCCAGTGTGGTTTG |
| *KLF9* | Forward | TGGCTGTGGGAAAGTCTATG |
|  | Reverse | GTCTGAGCGGGAGAACTTTT |
| *SOX4* | Forward | AGCGACAAGATCCCTTTCATTC |
|  | Reverse | CGTTGCCGGACTTCACCTT |
| *TP63* | Forward | TTCGGACAGTACAAAGAACGG |
|  | Reverse | GCATTTCATAAGTCTCACGGC |
| *ZNF185* | Forward | GGCTACAAGATGACCACTGAG |
|  | Reverse | CTCTGACCTCCGTTTCTGTTC |
| *ZNF853* | Forward | GTGCTGGAACTTCAATGTCTTG |
|  | Reverse | CTCTCCTGAGGCTCTTCCT |

**Global Gene Expression Analysis (RNA-sequencing)**

RNA from A549 cells treated with 5 mM MMA for 3 or 10 days was isolated as described above. Total RNA was sent to Active Motif for further processing and RNA-seq analysis. Briefly, RNA quality was assessed by BioAnalyzer, and only RNAs with RIN values between 8.7 and 10.0 were used. RNA-sequencing libraries were prepared using the Illumina TruSeq RNA Sample Preparation v2 Guide (Illumina Part # 15026495). Polyadenylated RNA was enriched from 1 µg total RNA. Libraries were sequenced on Illumina NextSeq 500 as paired-end 42-nt reads, to a depth of 40.2 – 54.7 M read pairs**. The “TopHat” algorithm was used to align the reads to the hg38 genome. The alignments (37.5-51.4 M aligned pairs) in the BAM files were further analyzed using the Cufflinks suite of programs (running consecutively: Cufflinks 🡪 Cuffcompare 🡪 Cuffdiff). Cufflinks was run using the hg38-genes as a reference data base. The cufflinks outputs were compared using cuffdiff.** The accession number for the raw sequencing data reported in this paper is GEO: GSE127001. A volcano plot visualizing the **genes that were significantly changed ≥1.5 fold** was created using EnhancedVolcano^10^. The top 100 significantly changed genes were log-tranformed and clustered using HierarchicalClustering on GenePattern^11^ using Euclidean distance and pairwise centroid-linkage, and was row centered. F**unctional annotation analysis on genes that were significantly changed ≥1.5 fold was performed using** DAVID: Database for Annotation, Visualization, and Integrated Discovery**^12^.** Gene sets based on annotations from the Gene Ontology database (http://www.geneontology.org) were used. **O**nly gene sets with 10 genes or more, and EASE Score, a modified Fisher Exact p-values, of 0.05 or less were evaluated.

**mRNA Overlap Analysis**

The overlap of significantly altered mRNAs between A549 cells treated with MMA for 10 days (≥1.5 fold change) and the significantly altered genes induced by SOX4 induction (≥2 fold change)^13^ was evaluated using the GeneOverlap algorithm^14^. F**unctional annotation analysis on the set of overlapping genes was performed using** DAVID: Database for Annotation, Visualization, and Integrated Discovery**^12^.** Gene sets based on annotations from the Gene Ontology database (http://www.geneontology.org) were used. **Gene sets with an** EASE Score, a modified Fisher Exact p-values, of 0.05 or less were evaluated.

**Lung Colonization Assay in Mice**

MDA-MB-231-lucifease cells were treated with human serum from young and old donors as described above prior to tail vein injection and lung colonization evaluation. To test the effect of MMA in lung colonization, luciferase-expressing MDA-MB-231 cells were treated with 1 or 5 mM MMA for 5 days. Similarly, SOX4 silenced MDA-MB-231-luciferase cells were also treated with 5 mM MMA for 5 days or with human serum from old donors, after which they were injected into the tail vein and lung colonization was evaluated as described before^2,15^. Briefly, female nu/nu athymic mice were injected with 100,000 cells in 100 μL PBS through tail vein injections. Metastases were monitored using IVIS Spectrum CT Pre-Clinical In Vivo Imaging System (Perkin-Elmer). 7-10 mice were used in each experimental group. After 6 weeks, luminescence was measured and quantified using the Living Image Software (Perkin-Elmer) to determine lung colonization. All animal studies followed the guidelines of and were approved by the Weill Cornell Medicine Institutional Animal Care and Use Committee.

**Orthotopic Xenograft Experiments in Mice**

To achieve increased circulatory MMA concentrations female nu/nu athymic mice were treated as described before^16^ with a dose escalation of a “low” (100 μg MMA/g mouse followed by 200 μg MMA/g mouse) or “high” MMA (200 μg MMA/g mouse followed by 400 μg MMA/g mouse) either by subcutaneous injections or through the drinking water. MMA treatments through either method did not cause behavioral changes or changes in drinking and feeding habits, nor did they cause changes in body weight throughout the duration of the experiments. Briefly, mice were treated for 16 days with the lower dose of MMA for both concentration groups after which the dose was doubled until the end of the experiment. For subcutaneous injections MMA (Sigma) was dissolved in 0.9% NaCl, adjusted to pH 7.4 with 6 M NaOH, and injections were performed daily during the time of the experiment. For delivery of MMA in the drinking water, MMA (Sigma) was dissolved in acidified water (as is standard IACUC procedure for mice husbandry) and it was replaced with a fresh solution every 3.5 days. Tumors were established in the mice after the first 8 days of MMA treatment by injection of 2 x 10^6 MDA-MB-231-luciferase cells in 100 μl 50:50 PSB and Matrigel (Corning) into the third mammary fat pad on the right side of each mouse. Primary tumors and metastases were monitored using IVIS Spectrum CT Pre-Clinical In Vivo Imaging System (Perkin-Elmer). To visualize the metastatic spread the primary tumors were covered and the upper body was imaged. The experiment continued until mice showed signs of significant illness or the primary tumors reached 10% of their weight at which point they were euthanized as specified by IACUC. At the time of euthanasia blood was collected to measure the serum MMA levels and primary tumor tissue was harvested. Time of natural death or euthanasia was used to create the Kaplan-Meier curves. All animal studies followed the guidelines of and were approved by the Weill Cornell Medicine Institutional Animal Care and Use Committee. Frozen tumor tissue was powderized with a mortar and pestle over liquid nitrogen. RNA was isolated from the powdered tissue and the qPCR was performed as described in the “gene expression analysis” section. For protein isolation powderized tissue was lysed with RIPA buffer for 30 minutes on ice and homogenized through a 22 gauge needle before centrifuged at full speed for 10 minutes. The proteins present in the lysate supernatants were then quantified and processed as described under “immunoblots for total cell lysates”.

**Measurement of MMA absolute concentration**

For intracellular measurements of MMA concentrations, metabolites were extracted and processed as described in the targeted metabolomics section. For mouse sera, metabolite extraction was performed in a mixture of ice/dry ice as previously described^17,18^. Briefly, 10 µL of plasma were extracted with 800 µL of 62.5% methanol containing 0.6 µg/ml of glutaric acid, followed by an addition of 500 µL of precooled chloroform. Samples were vortexed for 10 min at 4 °C and then centrifuged for other 10 min (max. speed, 4°C). After centrifugation, metabolites were separated in two phases divided by a protein layer: polar metabolites in the methanol/water (upper) phase and the lipid fraction in the chloroform (lower) phase. The samples were derivatized and measured as described before^19^. Briefly, polar metabolites were derivatized for 90 min at 37°C in 15 µL of 20 mg/ml methoxyamine in pyridine per sample. Subsequently, 15 µL of N-(tert-butyldimethylsilyl)-N-methyl-trifluoroacetamide, with 1% tert-butyldimethylchlorosilane were added to 7.5 µL of each derivative and incubated for 60 min at 60 °C. Polar metabolite fractions containing methylmalonic acid were dried at 4°C in a vacuum concentrator. Methylmalonic acid concentrations were analyzed by gas chromatography (7890A GC system) coupled to mass spectrometry (5975C Inert MS system) from Agilent Technologies. Metabolites were separated with a DB35MS column (30 m, 0.25mm, 0.25 µm) using a carrier gas flow of helium fixed at 1 ml/min. A volume of 1 µL of sample was injected with a split ratio 1 to 3 with an inlet temperature set at 270°C. For the detection of polar metabolites, the gradient was set at 100°C for 1 min ramped to 105°C at 2.5°C/min, then to 240°C at 3.5°C/min and finally to 320°C at 22°C/min. For the measurement of metabolites by mass spectrometry, the temperatures of the quadrupole and the source were set at 150°C and 230°C, respectively. An electron impact ionization energy fixed at 70 eV was applied and scan mode was used for the measurement of polar metabolites ranging from 100 to 600 a.m.u (mass). After the acquisition by GC-MS, an in house Matlab M-file was used to extract mass distribution vectors and integrated raw ion chromatograms. The natural isotopes distribution were also corrected using the method developed by Fernandez et al, 1996^20^. Peak areas were normalized to those of the internal standard glutaric acid.

**Statistical Analysis**

Data analyses were performed using Microsoft Excel and GraphPad Prism7. A two-tailed paired Student’s t test was used to determine significance when two conditions were compared; for experiments with more than two conditions a one-way ANOVA was used to determine significance. In both types of statistical analysis values of *p* < 0.05 were considered significant. Data are represented as the mean ± SEM (standard error of the mean), or individual data points and the mean ± SEM of at least three independent experiments performed. Number of replicates and animals are reported in the figure legends. For all experiments similar variances between groups were observed. Normal distribution of samples was not determined. In the DAVID functional annotation analyses for the RNA-seq experiments EASE Score, a modified Fisher Exact p-values, were used to determine significance as recommended.

**Data Availability**

RNA-seq data that support the findings of this study have been deposited in the Gene Expression Omnibus (GEO) under the accession code GSE127001, as well as in Extended Data Table 2. Source data information for the metabolomics experiment can be found in Extended Data Table 1. All raw data files, peak lists, and the sequence database for the proteomics analysis have been deposited in the MASSive repository (https://massive.ucsd.edu, ID#:MSV000084974) and are available for download at ftp://massive.ucsd.edu/MSV000084974. All the raw images for the western blots as well as all other data supporting the findings of this study are available from the corresponding authors upon reasonable request.

**Code Availability**

Code information for quantification of transwell migration and invasion assays is available from the corresponding authors upon reasonable request.
